# Ribosomal protein RACK1 facilitates efficient translation of poliovirus and other viral IRESs

**DOI:** 10.1101/659185

**Authors:** Ethan LaFontaine, Clare M. Miller, Natasha Permaul, Alex G. Johnson, Elliot T. Martin, Gabriele Fuchs

## Abstract

Viruses have evolved various strategies to ensure efficient translation using host cell ribosomes and translation factors. In addition to cleaving translation initiation factors required for host cell translation, poliovirus (PV) uses an internal ribosome entry site (IRES) to bypass the need for these translation initiation factors. Recent studies also suggest that viruses have evolved to exploit specific ribosomal proteins to enhance translation of their viral proteins. The ribosomal protein receptor for activated C kinase 1 (RACK1), a protein of the 40S ribosomal subunit, was previously shown to mediate translation of the 5′ cricket paralysis virus and hepatitis C virus IRESs. Here we found that while translation of a PV dual-luciferase reporter shows only a moderate dependence on RACK1, PV translation in the context of a viral infection is drastically reduced. We observed significantly reduced poliovirus plaque size and a delayed host cell translational shut-off suggesting that loss of RACK1 increases the length of the virus life cycle. Our findings further illustrate the involvement of the cellular translational machinery in PV infection and how viruses usurp the function of specific ribosomal proteins.

## Introduction

Since viruses rely on cellular translation factors and ribosomes for translation of viral proteins, viruses and host cells battle for these critical resources. Viral double-stranded RNA activates interferon-induced, double-stranded RNA-activated protein kinase (PKR), which phosphorylates translation initiation factor eIF2α leading to inhibition of viral and cellular translation (1–3). To prevent eIF2α phosphorylation and translational shut-off, viruses target PKR. Some viral proteins directly bind to PKR to prevent its activity, other viruses degrade PKR or alter its subcellular localization (4–9). To efficiently compete for ribosomes, many viruses use translation initiation mechanisms distinct from cellular mRNA translation initiation, which uses canonical cap-dependent translation. All cellular mRNAs are transcribed in the nucleus, where they are also capped and polyadenylated. After export into the cytoplasm, the 5′ m^7^GpppN cap is bound by the cap binding protein eukaryotic initiation factor 4E (eIF4E) and the polyA-tail is bound by the polyA binding protein (PABP). Through binding of the scaffolding protein eIF4G to eIF4E and PABP, the mRNA is circularized. With help of the RNA helicase eIF4A, the 40S ribosomal subunit in complex with eIF2 and eIF3 scans the 5′ untranslated region (UTR) in an ATP-dependent manner until the start codon is reached and recognized. After 60S ribosomal subunit joining and GTP hydrolysis by eIF5B elongation can proceed. To prevent cap-dependent translation poliovirus (PV) and other viruses of the *Picornaviridae* family target these eIFs. Specifically, PV proteases 2A and 3C cleave eIF4G, and PABP and eIF5B, respectively (10–16). Cleavage of these essential translation factors shuts-off host cell translation, while PV uses an internal ribosome entry site (IRES) for translation of the viral polyprotein that does not rely on these translation factors (16, 17). Viruses not only prevent global translation inhibition in the host cell, they also employ strategies that specifically decrease translation of cellular mRNAs.

In addition to targeting translation initiation factors, several viruses have shown direct usage of ribosomal proteins to increase their viral translation. Lee et al. performed an siRNA screen and identified eight ribosomal proteins including eL40, that are not required for cell viability, but negatively affect Vesicular Stomatitis Virus (VSV) and other related viruses (18). Ribosomal protein eL40 was dispensable for viruses that use IRES-mediated translation, but regulated a subset of cellular mRNAs with diverse functions (18). In contrast to eL40, eS25, a protein located near the head of the 40S ribosomal subunit, mediates translation of viruses that initiate translation using IRESs (19, 20). eS25 directly interacts with the hepatitis C virus (HCV) and the intergenic (IGR) cricket paralysis virus (CrPV) IRESs in cryo-EM structures (21–24) and is required for high-affinity binding of the 40S ribosomal subunit to the CrPV IRES (19). Further, eS25 not only facilitates translation of other IRESs such as encephalomyocarditis virus (EMCV) and PV, but also regulates translation of cellular IRES-containing mRNAs (20). More recently, another ribosomal protein, receptor for activated C kinase 1 (RACK1) has been shown to be exploited by different viruses.

RACK1 is a core ribosomal protein (25) that belongs to the tryptophan-aspartate repeat (WD-repeat) protein family. The seven-bladed β-propeller structure of RACK1 is located near the mRNA exit tunnel where it makes contacts with the ribosomal RNA through lysine and arginine residues and neighboring ribosomal proteins (26–28). RACK1 is often termed a scaffolding protein and has been implicated in a variety of biological function on and off the ribosome. In addition to binding to its eponym protein kinase C βII (PKCβII) and being involved in cellular signaling via Src protein-tyrosine kinase (29–31), RACK1 has been shown to interact with the microRNA machinery (32), bind eIF6 to regulate the 60S ribosomal subunit (33) and regulate ribosome-associated quality control (34, 35). At the level of tissues and organisms, RACK1 regulates axonal growth (36), neural tube closure in *Xenopus laevis* (37), and is essential for development in mice (38), *Drosophila melanogaster* (39) and *Arabidopsis thaliana* (40, 41), but appears to be dispensable in single cell organisms such as yeast (27). Directly and/or indirectly, RACK1 also influences translation of cellular mRNAs. The *Saccharomyces cerevisiae* RACK1 homolog, Asc1, facilitates efficient translation of mRNAs containing a short open reading frame (42), while in mammalian cells, RACK1 appears to stimulate cell proliferation in a PKCβII-dependent manner (30, 43, 44).

At the intersection of cellular signaling and translational regulation, RACK1 represents a critical regulatory target for many viruses. Vaccinia virus, which belongs to the poxviruses and contains a dsDNA genome, but replicates exclusively in the cytoplasm, encodes a kinase that phosphorylates a flexible loop in RACK1 (45). Through phosphorylation, this RACK1 loop is now negatively charged, which allows for translation of the poxvirus polyA-leader containing mRNAs (45). In plants, where polyA-leader sequences are commonly found, this RACK1 loop contains several glutamic acid residues, hence poxvirus evolution likely rediscovered efficient translation of polyA-leaders through phosphorylation of RACK1. Viruses from the *Dicistroviridae* family encode two polyproteins, and translation of each polyprotein is mediated by an IRES (46). In contrast to eS25, RACK1 is dispensable for translation of the CrPV IGR IRES, but its loss inhibits the translation of both the 5′ IRES of CrPV as well as the HCV IRES (47).

The finding that RACK1 facilitates efficient translation of the HCV IRES prompted us to explore if the need for RACK1 is more broadly conserved. Using a RACK1 knockout cell line generated by CRISPR-Cas9 genome editing (45), we first tested translation using dual-luciferase constructs. We found that HCV, EMCV and PV IRES translation are all reduced in cells lacking RACK1. Although the effect on PV translation in the context of the dual-luciferase reporter is moderate, loss of RACK1 causes a significant decrease in the PV plaque size. This decrease is due to attenuated translation prior to and post translational shut-off, suggesting that the virus life cycle lengthened in cells lacking RACK1.

## Results

### RACK1-FLAG is incorporated into polysomes

To investigate the function of mammalian RACK1 in translation, we established a functional rescue by expressing RACK1-FLAG in HAP1-derived CRISPR genome edited RACK1 knockout cell line RACK1 KO #1 described by Jha et al. (45) using lentiviral transduction (48). HAP1 cells are a near-haploid human cell line derived from chronic myelogenous leukemia KBM-7 cells (49). RACK1 was undetectable in RACK1 KO #1 and RACK1 KO #2 cell lines. Following lentiviral transduction of RACK1-FLAG into RACK1 KO #1 cells, RACK1 levels were partially restored (figure 1A). To examine incorporation of FLAG-tagged RACK1 into translating ribosomes rather than other high molecular weight cytosolic complexes, we performed polysome analysis by sucrose gradient ultracentrifugation. When cell lysate is treated with the translation elongation inhibitor cycloheximide, translation will be stalled. Upon sucrose gradient ultracentrifugation, the translating ribosomes, polysomes, are separated from the ribosomal subunits. When sucrose gradient analysis was performed on wildtype HAP1 and RACK1-FLAG expressing RACK1 KO #1 cells, no major differences in the overall polysome profile were detected (figures 1B and 1C).

**Figure 1:**
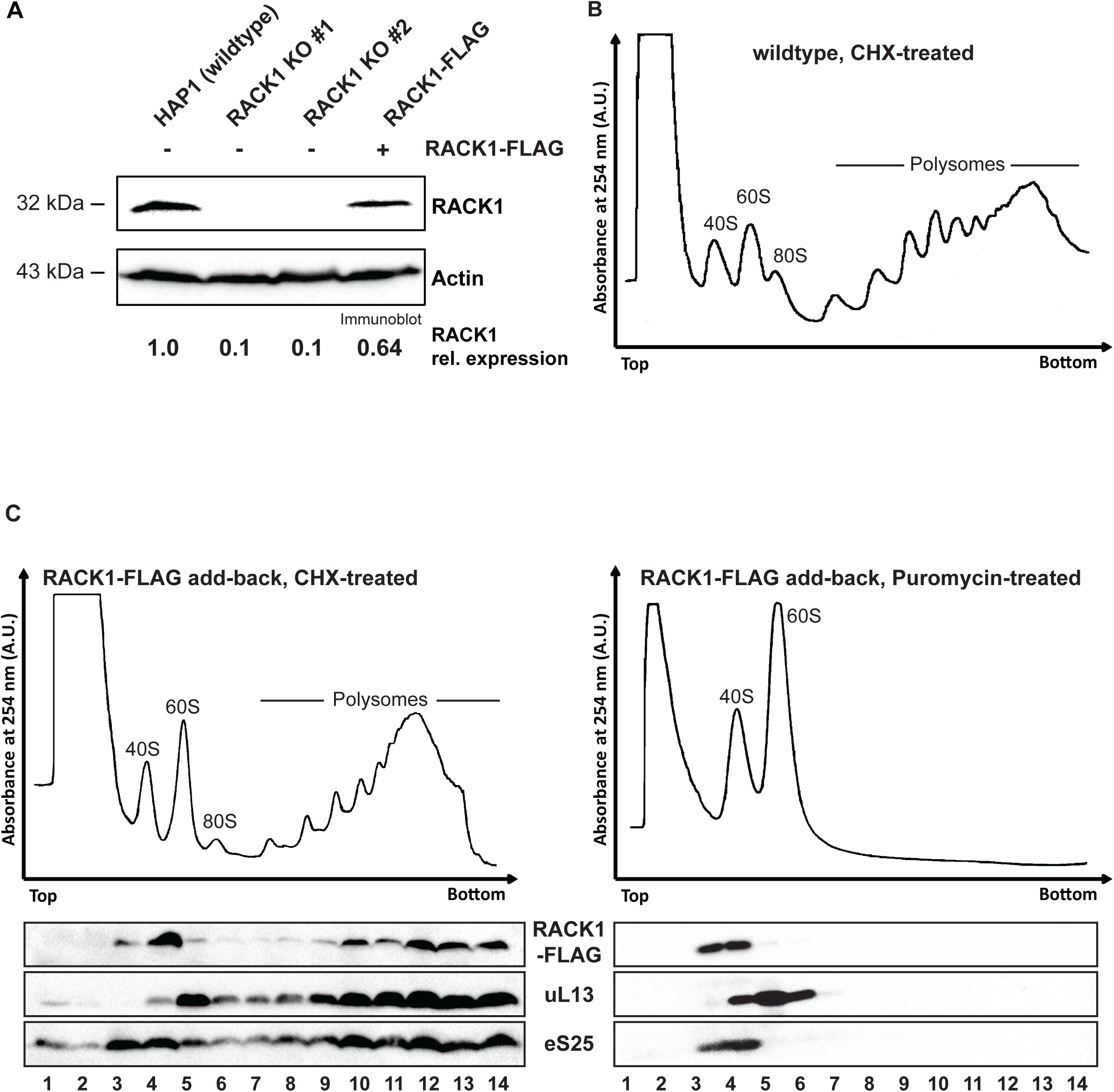
RACK1-FLAG is incorporated into polysomes. (A) RACK1 levels can be partially restored in RACK1 KO cells by expression of RACK1-FLAG. RACK1 levels in the different cell lines were quantified on the immunoblot analysis of RACK1 and the loading control actin. (B) Polysome trace of HAP1 wildtype cells. Cell lysate was separated in 10-50% sucrose gradient and absorbance at 254 nm was measured. (C) Sucrose gradient analysis of FLAG-tagged RACK1 protein. In cells treated with the translation elongation inhibitor cycloheximide, RACK1-FLAG detected by immunoblotting using an anti-FLAG antibody co-sediments with polysomes (fractions 9-14). Upon treatment of cell lysate with the translation elongation puromycin, which separates actively translating ribosomes into the ribosomal subunits, RACK1-FLAG co-sediments with 40S ribosomal subunits in lighter sucrose gradient fractions (fractions 3-4). Immunoblot analysis for eS25 and ul13 are used as indicators for sedimentation of protein components of the small and large ribosomal subunits, respectively.

We found that RACK1-FLAG co-sedimented in the polysomal fractions 9-14 (figure 1C, left panel) with eS25 and eL13a, ribosomal proteins of the 40S and 60S ribosomal subunits, respectively. Although this result suggested that RACK1-FLAG was incorporated into polysomes, we could not exclude that it sedimented in heavy sucrose fractions because it formed aggregates. To further validate that RACK1-FLAG indeed was incorporated into polysomes, we treated cell lysate with puromycin. Puromycin is a tRNA analog, which stalls translation elongation and releases the growing peptide chain. When cell lysate treated with puromycin is heated to 37°C, the two ribosomal subunits are separated, and the mRNA is released (50). Puromycin treatment alters the sedimentation pattern for ribosomal proteins, which now sediment in lighter sucrose fractions where the ribosomal protein subunits are found. Following puromycin treatment, RACK1-FLAG now sedimented in the lighter sucrose gradient fractions 3 and 4 (figure 1C, right panel), where it again co-sediments with eS25. Taken together, these results indicate that RACK1-FLAG is incorporated into translating ribosomes and likely fully functional.

### RACK1 mediates translation of viral IRESs

Loss of RACK1 has been previously shown to inhibit translation of the HCV and CrPV 5′ IRES (47) raising the possibility that RACK1 generally facilitates viral IRES-mediated translation. To test this hypothesis, we used dicistronic luciferase reporters, in which translation of the *Renilla* luciferase uses canonical cap-dependent translation initiation, while translation of the Firefly luciferase is mediated by a viral IRES (figure 2A). We tested the importance of RACK1 for translation of four viral IRESs, specifically PV, EMCV, HCV and CrPV intergenic IRESs. These IRESs represent four major types of viral IRESs and use different mechanisms for translation initiation (4). None of these viral IRESs use the cap-binding function of eIF4E, although a recent study showed that eIF4E stimulates the helicase activity of eIF4A on the PV IRES independent of its cap-binding function (52). In contrast to PV, neither the EMCV nor the HCV IRES use a scanning mechanism but instead directly recruit the ribosome to the start codon (53). While the EMCV IRES requires all canonical translation initiation factors except for eIF4E, translation initiation of the HCV IRES uses a more limited set of translation initiation factors. In agreement with previous observations, loss of RACK1 did not alter translation of the CrPV intergenic (IGR) IRES, but inhibited translation of the HCV IRES (figures 2B and 2C, and (47)). In addition, we observed that RACK1 also facilitated translation of the EMCV and PV IRESs (figures 2B and 2C). Expression of RACK1-FLAG in RACK1 knockout cells partially rescued the defect in IRES-mediated translation (figure 2C). The observed rescue approximately corresponded to the expression level of RACK1-FLAG (figures 2C and 1A). Taken together, these data support the need for RACK1 to facilitate translation of HCV, EMCV and PV IRESs, but not CrPV IGR IRES.

**Figure 2:**
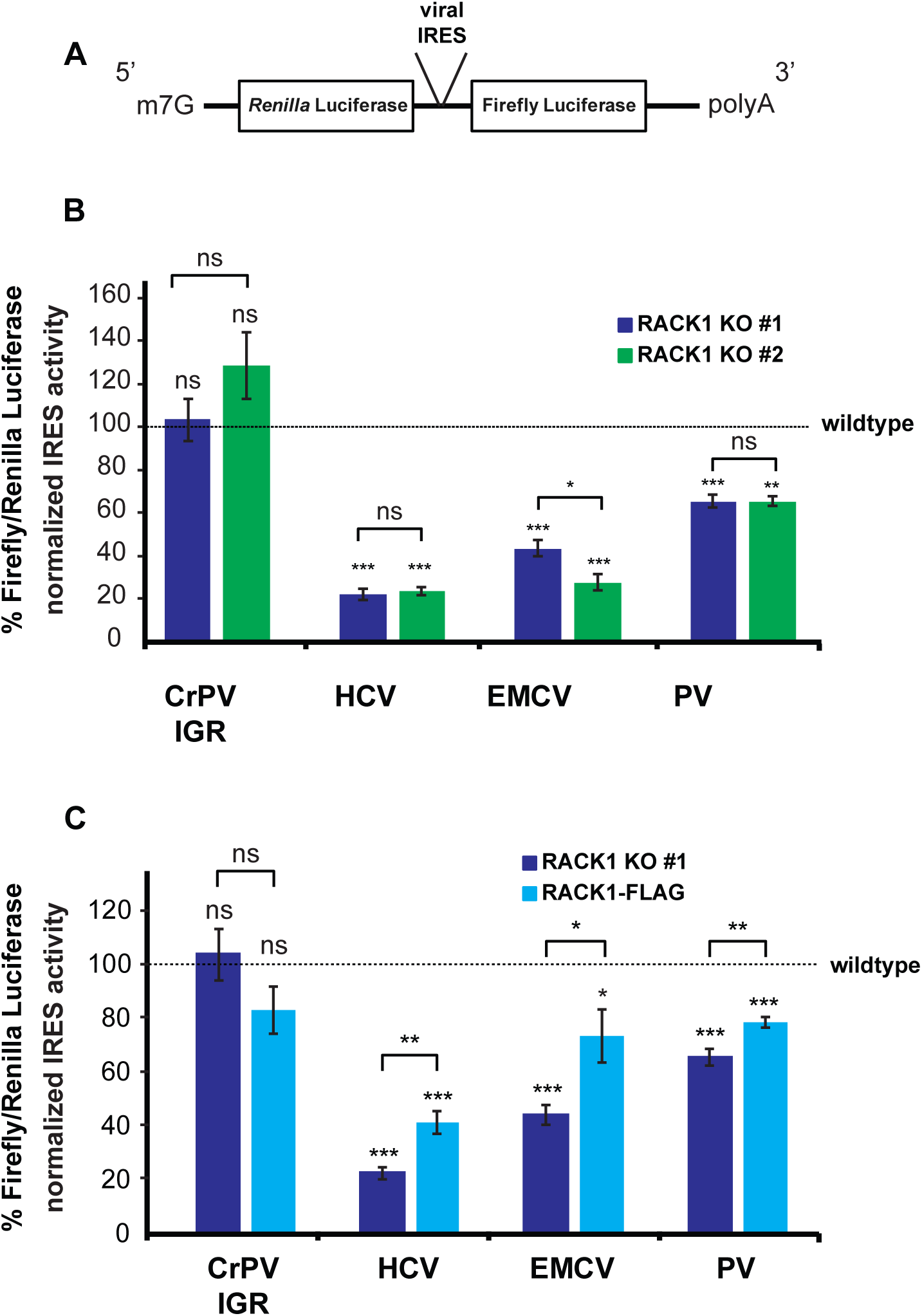
RACK1 facilitates translation of viral IRESs. (A). Schematic overview of dual luciferase construct used in assays. The *Renilla* luciferase open reading frame is translated via m7G cap-dependent translation, while the viral IRES located in the intergenic region between the two coding sequences mediates translation of the Firefly luciferase. (B) Translation efficiency of CrPV, HCV, PV and EMCV dual luciferase reporters in both RACK1 KO cells normalized to HAP1 cells (dotted line). Error bars represent the standard error of the mean of at least three independent experiments. ** p-value < 0.01; *** p-value < 0.001 (C) Translation of HCV, EMCV and PV IRESs correlates with RACK1 levels, but translation of CrPV IGR IRES is RACK1 independent. Ratios of Firefly/*Renilla* normalized to HAP1 ratios (dotted line). Error bars represent the standard error of the mean of at least three independent experiments. * p-value < 0.05; ** p-value < 0.01; *** p-value < 0.001

### PV plaque diameters are decreased in cells lacking RACK1

To test if the reduction of PV IRES-mediated translation impacts the virus during infection, we performed PV plaque assays in wildtype, RACK1 KO #1, and RACK1-FLAG add-back cells and measured both PV plaque diameter and plaque numbers. Following infection with the Mahoney strain of PV, we observed a significant decrease in the PV plaque size in cells lacking RACK1 as compared to wildtype and RACK1-FLAG add-back cells (figure 3A). In contrast, the number of PV plaques was not significantly altered in any of the cell lines (figure 3B). Together, these data indicate that infectious particles are similarly efficient at establishing an infection independent of cellular RACK1 levels, however, the infectious cycle and virus spread may be impaired.

**Figure 3:**
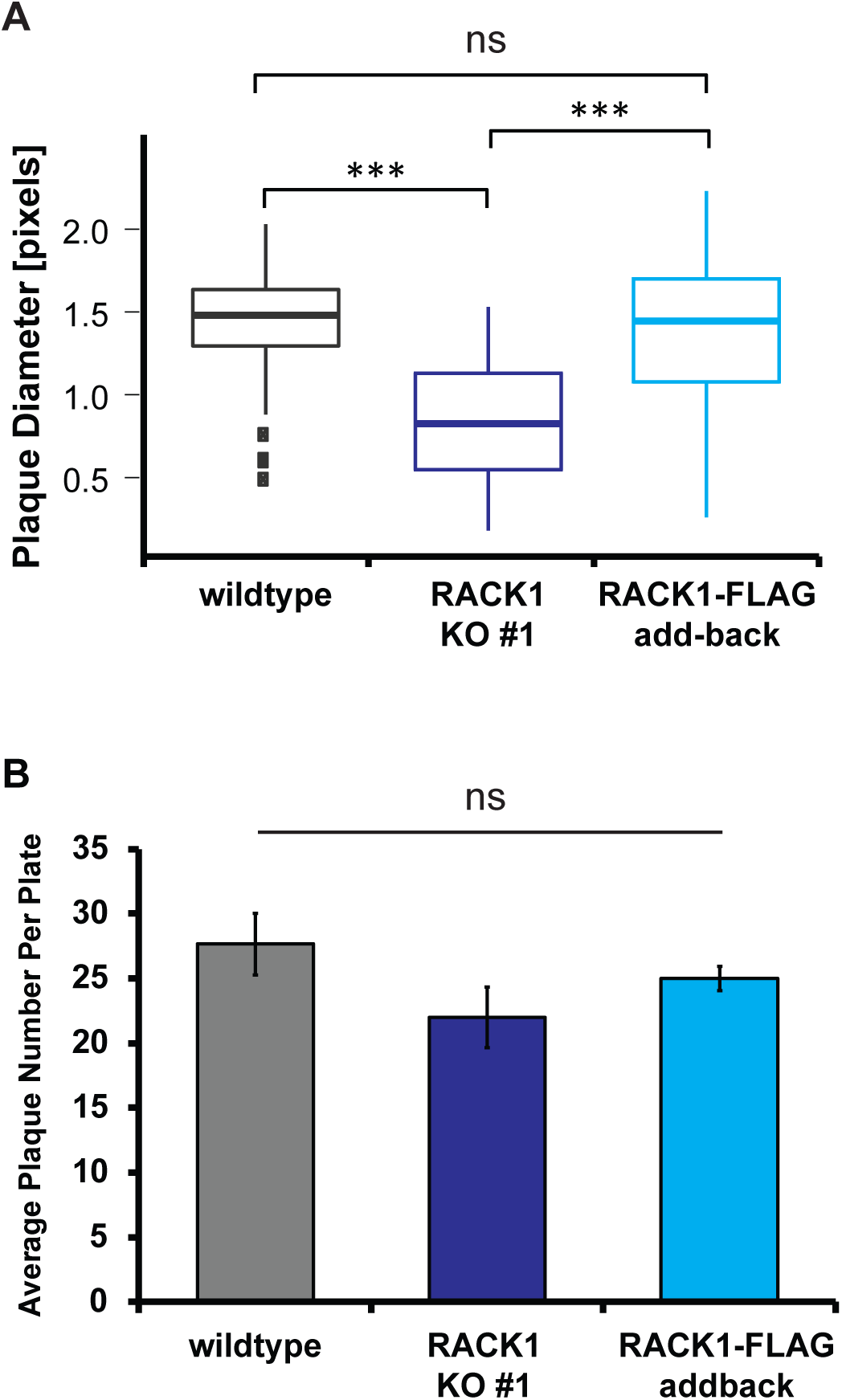
PV plaque size is reduced in cells lacking RACK1. (A) Analysis of PV plaque assays. PV plaques in RACK1 KO #1 cells are smaller compared to PV plaques in wildtype and RACK1-FLAG addback cells (left panel). Cells were infected at identical MOI; 42 h post infection, cells were stained with crystal violet and poliovirus plaque diameters were measured. Error bars represent the standard deviation within the population of three independent experiments. The box indicates the data between the interquartile range (IQR) between the 25^th^ and 75^th^ percentile, the median is indicated by the thick line within the box. The thin vertical bars represent the minimum and maximum data points (Q1-1.5*IQR, Q3+1.5*IQR). Outliers in the wildtype cells are indicated by filled squares. (B) The number of visible poliovirus plaques does not differ between wildtype, RACK1 KO #1 and RACK1-FLAG cell lines. Cells were infected at identical MOI; 42 h post infection, cells were stained with crystal violet and poliovirus plaques were counted. Error bars represent the standard error of the mean of at least three independent experiments.

### Loss of RACK1 impairs PV translation during the entire virus life cycle

Upon PV infection, the PV genome must be translated to give rise to the viral proteins, which include the viral proteases 2A and 3C. When levels of 2A and 3C are sufficiently high, these viral proteases cleave translation factors eIF4G and PABP, which shuts off translation in the host cell. Our observation that PV plaques are reduced in cells lacking RACK1 made us wonder whether loss of RACK1 prolonged the viral life cycle and delayed host-cell translation shut-off. To test whether translational shut-off was delayed in RACK1 KO cells, we performed ^35^S pulse-labeling experiments in mock-infected and PV-infected cells at a multiplicity of infection (MOI) of 1. When newly synthesized proteins were metabolically labeled 3 hours 5 minutes post infection and lysates revolved by SDS-PAGE we observed that wildtype and RACK1-FLAG add-back cells started to show the characteristic PV banding pattern observed in the positive control of cells infected at MOI = 10 and harvested at the same time (figure 4A) (16). In contrast, the protein pattern in the RACK1 KO cell lines was similar to the pattern in the mock-infected cells, in that cellular proteins were metabolically labeled, and viral proteins are absent. These data indicate that the time until PV-induced translational shut-off of the host cell was indeed delayed (figure 4A). We next asked if PV translation also remained at lower levels in RACK1 KO cells post translational shut-off. To address this question, we monitored translation of a PV-Luciferase replicon (PV-Luc) during the initial phase of translation and translation during viral replication. Instead of the viral structural proteins, the PV-Luc replicon encodes a luciferase open reading frame fused to the PV non-structural proteins.

**Figure 4:**
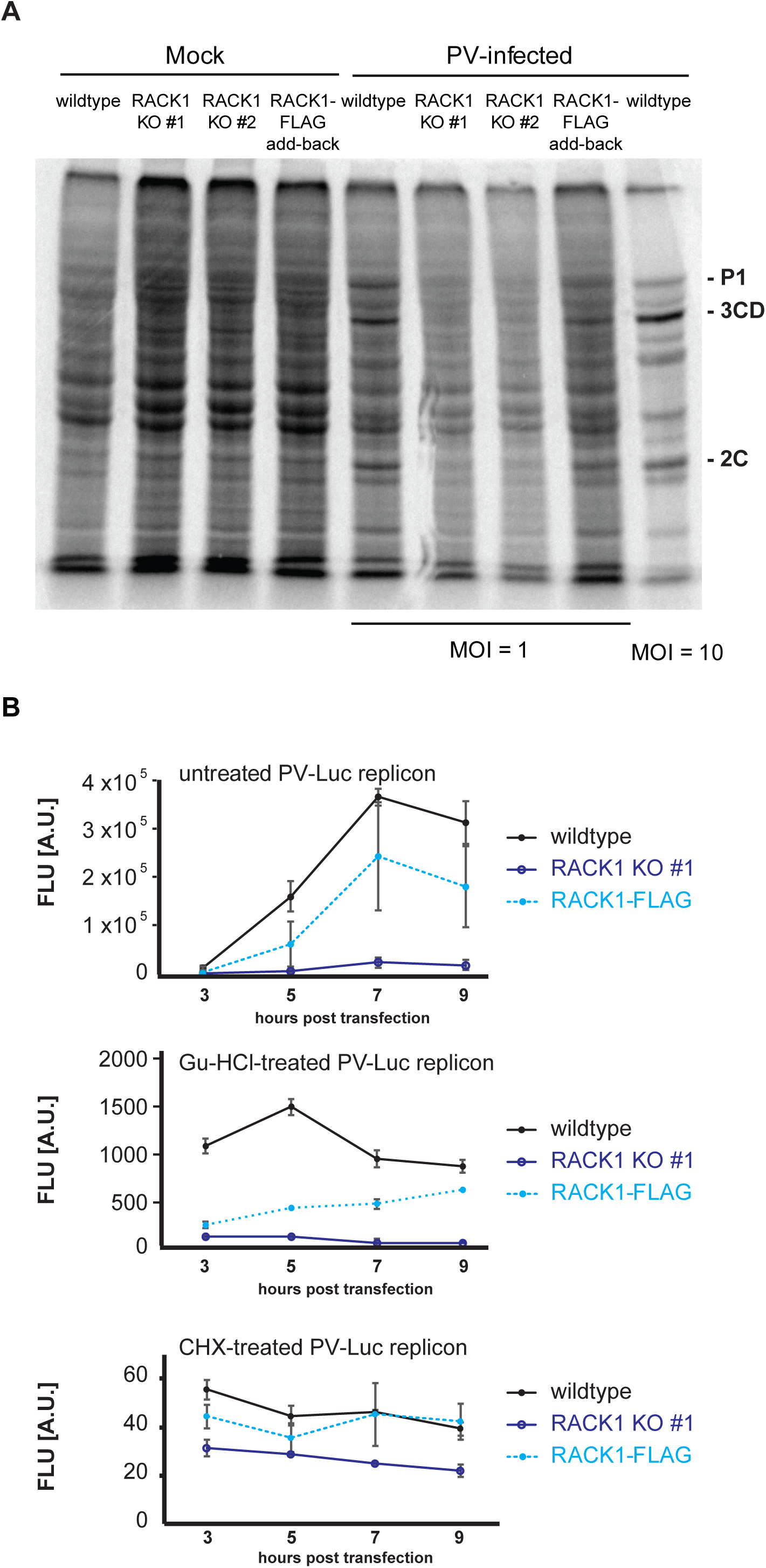
RACK1 is required for efficient PV translation prior and post host cell translational shut-off. (A) ^35^S metabolic pulse labeling of uninfected (mock) and PV-infected cells. Cells were mock-infected or infected with PV Mahoney at MOI = 1 (MOI = 10 for positive control) and ^35^S pulse-labeled for 10 min at 3 hours and 5 min post infection. Total protein lysates were separated in 10% SDS-PAGE and visualized by exposure to a phosphor-screen. Poliovirus-specific protein products P1, 3CD, and 2C are indicated (72). (B) Expression of a PV-Luc replicon is inefficient in cells lacking RACK1. HAP1, RACK1 KO #1, and RACK-1 FLAG cells were transfected with *in vitro* transcribed PV-Luc replicon RNA. An aliquot of cells was removed 3, 5, 7, and 9 hours post RNA transfection and Firefly luminescence was measured. Error bars represent the standard error of the mean of at least three independent experiments. Top panel: PV-Luc translation is decreased in cells lacking RACK1 during the replication phase. Middle panel: To limit the observation to the early translation phase, PV-Luc replication was inhibited with 2 mM guanidine hydrochloride immediately after RNA transfection. Bottom panel: Translation of all PV-Luc reporters is completely inhibited upon treatment with 25 μg/ml cycloheximide. Cells were treated with cycloheximide immediately after RNA transfection.

Early during infection, luciferase measurements reveal initial translation of the replicon. Once negative strand synthesis has occurred, the PV genome will start replicating, which will result in greater translation of the replicon and largely increased luciferase production. We transfected *in vitro* transcribed PV-Luc RNA into wildtype, RACK1 KO #1 and RACK1-FLAG cells, harvested protein lysates 3, 5, 7, and 9 hours post transfection and measured luciferase levels by luminescence. As expected, we observed robust luciferase production in wildtype cells, while levels of luciferase in RACK1 KO #1 remained more than 10-fold lower (figure 4B, top panel). Consistent with our previous results, RACK1-FLAG expression in the KO cell line partially rescued luciferase expression. To help distinguish viral translation and replication stages, we immediately treated cells with guanidine hydrochloride, a drug that inhibits viral replication (54). Thus, the measured amount of luciferase produced in these experiments only represents protein production prior to viral replication. Although luciferase levels in RACK1 KO#1 and RACK1-FLAG cells were comparable at 3 hours post transfection, luciferase levels in the RACK1-FLAG cell line increased over time, reaching levels comparable to luciferase levels in wildtype cells, while cells lacking RACK1 KO#1 failed to accumulate significant levels of luciferase (figure 4B, middle panel). When cells were treated with the translation inhibitor cycloheximide immediately after PV-Luc reporter transfection, luciferase levels measured in all cells were at background levels (figure 4B, lower panel), indicating that translation of the viral genome had been completely blocked.

Taken together, this data indicates that RACK1 not only mediates translation of the HCV IRES but is also critical for efficient translation of the EMCV and PV IRESs using dicistronic luciferase assays. Cells lacking RACK1 also show a reduced PV plaque size, likely due to inefficient PV translation prior to as well as post host cell translational shut-off, which further extends the PV life cycle.

## Discussion

The ribosomal protein RACK1 interacts with numerous cellular proteins and has been thought to function as a scaffolding protein that connects cellular signaling pathways with the ribosome and the translation machinery (55). In addition, it has been previously shown that RACK1 is important for translation of the HCV IRES (47). Although HCV, PV and EMCV all use IRESs for translation initiation, the PV and EMCV IRESs rely on more translation factors than the HCV IRES. These translation factors could compensate for the function of RACK1, which prompted us to test whether RACK1 is also needed for translation of other viral IRESs. We thus employed RACK1 knockout cells generated by CRISPR-Cas9 mediated genome editing (45) and transduced them with lentiviruses to express RACK1-FLAG, which was incorporated into translating ribosomes (figure 1). In contrast to others who have reported that RACK1 depletion reduces cap-dependent translation (30, 56), we do not observe major changes in cap-dependent translation in RACK1 KO cells ((45), figure 4A and unpublished data). It is possible that the contrasting observations are due to cell line specific differences. While HAP1 cells are derived from the chronic myelogenous leukemia (CML) cell line KBM-7, all studies that found that RACK1 influenced cap-dependent and -independent translation were performed in HEK293, HEK293T, and HeLa cells (30, 31, 56). Since RACK1 stimulates global translation in a PKCβII-dependent manner (30, 43), potentially higher PKCβII expression levels found in HEK293 and HeLa cells or other cell line-specific differences might explain the opposing effect on translation ((57), https://www.proteinatlas.org/ENSG00000166501-PRKCB/cell).

To test the involvement of RACK1 in IRES-mediated translation beyond CrPV and HCV IRESs, we used dicistronic luciferase constructs (figure 2A) containing the EMCV and PV IRESs, two other well-characterized viral IRESs. We found that RACK1 not only facilitates translation of the HCV IRES, but also of the EMCV and PV IRESs (figure 2). In agreement with previous work by Majzoub et al., translation of the CrPV IGR IRES occurred in a RACK-independent manner (47). To examine if RACK1 also plays a critical role during PV infection, we next infected wildtype, RACK1 KO and RACK1-FLAG expressing cells with the Mahoney strain of PV and performed plaque assays. While loss of RACK1 caused significantly smaller PV plaques (figure 3A), almost all infectious particles are able to start a successful infection (figure 3B). This finding indicates that RACK1 is partially dispensable for PV IRES-mediated translation, but also suggests that RACK1 specifically influences the translation efficiency of the PV IRES-containing RNA. Several groups have found RACK1 to regulate autophagy by directly interacting with Atg5 (58) and by enhancing Atg14L-Beclin 1-Vps34-Vps15 complex formation (59). However, neither Atg5 nor Beclin 1 impact PV proliferation (60) making it unlikely that the observed phenotype is due to changes in the autophagy pathway.

Using metabolic labeling and translation of a PV-Luc reporter we further showed that PV translation is attenuated in cells lacking RACK1 (figure 4) both pre and post translational shut-off (figure 4B). Decreased PV translation early during the virus life cycle lengthens the time until critical quantities of poliovirus proteases 2A and 3C are produced. These two proteases cleave translation factors eIF4G and PABP, respectively, cause host cell translation shut-off and enable viral replication (11, 14, 61). In cells lacking RACK1, translational shut-off of the host cell takes longer compared to RACK1-expressing cells (figure 4A). Further, RACK1 is not only critical prior to translational shut-off, but our PV-Luc reporter assay also showed that cells lacking RACK1 do not efficiently replicate the PV genome, while cells expressing RACK1 start replication 3 h post transfection (figure 4B, top panel). Although RACK1 is not essential for PV translation, it enhances the translation efficiency of PV and other viruses, indicating RACK1 acts as a pro-viral host protein.

Interestingly, our findings are reminiscent of ribosomal protein eS25, previously shown to mediate both viral and cellular IRES-mediated translation (19, 20). Both eS25 and RACK1 have a greater impact on HCV IRES translation, reducing IRES activity by about 75%, while translation of the PV IRES is less sensitive to eS25 and RACK1 levels. Curiously, the EMCV IRES appears to depend more on RACK1 than eS25, since loss of RACK1 causes a 60% decrease in IRES activity, while loss of eS25 only decreases EMCV IRES activity by 40%. Similarly, though, shRNA-mediated depletion of eS25 resulted in a 2-fold decrease in PV viral titers (20), which indicates that loss of eS25 might also lengthen the viral life cycle prior and post translational shut-off, however, no PV plaque size of a direct virus plaque assay was reported. In contrast to RACK1, eS25 reduction was found to inhibit all viral IRESs, including the CrPV IGR IRES (19). Both RACK1 and eS25 are ribosomal proteins that are usurped by viruses to enhance viral translation.

Our findings may be explained by three potential models for how RACK1 acts on PV translation. First, RACK1 could directly enhance the affinity of the ribosome for the PV IRES, for example by stabilizing IRES docking. Indeed, Landry et al. showed that the CrPV IRES is unable to bind to 40S ribosomal subunits lacking eS25 (19). Since the 40S ribosomal subunit is not directly recruited by the PV IRES but involves a scanning mechanism, *in vitro* ribosome affinity can only be measured in the presence of purified translation initiation factors, which is quite challenging. However, 40S ribosomal subunits lacking RACK1 directly bind the HCV IRES with an affinity similar to wildtype 40S ribosomal subunits (48). Further, ribosomes lacking RACK1 are also able to form 80S ribosomes at high concentrations of magnesium (48) indicating that RACK1 is neither involved in 40S nor 80S:HCV IRES complex formation. Although we cannot exclude a direct contribution of RACK1 to 40S binding or 80S complex formation with the EMCV and PV IRESs, we believe that the evidence for the HCV IRES suggests that RACK1 might employ a mechanism distinct from eS25.

Second, RACK1 might directly or indirectly affect the structure of the ribosome:PV IRES complex, for example by stabilizing a translation-favorable IRES conformation. Such structural rearrangement has been observed for the HCV IRES (24) where domain 2 of the HCV IRES, which interacts with eS25, undergoes a ∼55 Å movement. This movement switches the 40S ribosomal subunit from an open conformation for mRNA loading to a closed conformation with the initiator tRNA tightly bound to the P-site (24). Although domain II of the HCV IRES does not contribute to the binding affinity of the HCV IRES to the 40S ribosomal subunit (21, 62, 63), eS25 has been shown to have a critical function for HCV IRES translation (19, 23). The increased size of the PV IRES, which is almost double the size of the HCV IRES, and the complex translation initiation pathway of the PV IRES involving ribosome scanning and several more translation initiation factors has prevented the acquisition of a cryo-EM structure to test these models. Thus, in cell structure probing techniques such as selective 2′-hydroxyl acylation analyzed by primer extension (SHAPE) coupled with the replication inhibitor guanidine hydrochloride, may be required to provide valuable insights into the PV-IRES ribosome structure in the future (64).

Third, RACK1 is not near the HCV IRES binding interface as revealed by the cryo-EM structure of the HCV IRES:40S complex, further indicating that RACK1 unlikely affects IRES binding. In contrast to the CrPV IGR IRES, both the 5′ CrPV and HCV IRESs require eIF3, which has an extensive binding surface on the 40S ribosomal subunit. Translation initiation factor eIF3 is composed of 13 protein subunits and binds to a large surface of the 40S ribosomal subunit (65, 66). While several eIF3 subunits are essential for its function, other factors such as eIF3H and eIF3J are dispensable (47, 67). Of these dispensable functions, loss of eIF3J mimics the RACK1 loss-of-function phenotype on HCV and CrPV 5′ IRES translation (47), possibly by altering the conformation of the mRNA entry channel (48). Again, the lack of a cryo-EM structure of the large PV IRES bound to the 40S ribosomal subunits prevents us from using structural data to gain insights into the mechanism of RACK1 on PV translation. Instead, biological approaches such as crosslinking-immunoprecipitation (CLIP) on PV-infected cells will have to be used to identify potential interactions between RACK1 and the PV IRES (68).

Although it is unclear how RACK1 facilitates PV translation and if the way RACK1 is used by PV is identical to HCV translation, the similarities to eS25 roles in IRES-mediated translation are striking. While the two ribosomal proteins appear to use distinct mechanisms, both might alter the conformation of the mRNA entry channel and/or the transitioning between the open and closed 40S conformation (24, 47). These findings further raise the question which cellular RNAs are translationally regulated by RACK1 and eS25 and whether the mRNA targets are distinctly different. The observations that neither RACK1 nor eS25 are largely involved in canonical cap-dependent translation (19, 45) suggests that the specific cellular mRNAs relying on these proteins might be translationally highly regulated. Hertz et al. found that cellular IRES-containing mRNAs are less efficiently translated in eS25-depleted cells (20), indicating a role of eS25 in cap-independent translation. However, given the indirect interaction of RACK1 with the HCV IRES via eIF3, further studies will be needed to determine if RACK1 also facilitates cap-independent translation or if it acts a translational enhancer of specific mRNAs (69).

Ribosomal protein RACK1 not only enhances translation of HCV and the CrPV 5’ IRES, it also enhances translation of other viral IRES-containing RNAs such as EMCV and PV. PV-infected cells lacking RACK1 inefficiently translate the viral RNA, which lengthens the virus life cycle. These results suggest that targeting RACK1 could be used as an antiviral strategy, but more research into the cellular mRNAs that rely on RACK1 for translation is needed.

## Materials and Methods

### Cell culture

HAP1 cells purchased from Horizon USA (C859), HAP1-derived RACK1 knockout cell lines #1 and #2 (45), and RACK1-FLAG addback cell lines were cultured in Iscove’s modified Dulbecco’s medium (IMDM; Corning) supplemented with 5% fetal bovine serum and 2 mM L-glutamine. HEK293FT cells (Thermo Scientific) grown in DMEM (Gibco) supplemented with 5% fetal bovine serum and 2 mM L-glutamine were used to generate lentivirus particles for cellular transductions. Cultures were confirmed negative for mycoplasma using DAPI staining.

### Viral-Transduction of RACK1 Add-Back Cell Line

RACK1 cDNA was cloned with primers Forward 5′ATGACTGAGCAGATGACCCTTCG3’ and Reverse 5′CTAGCGTGTGCCAARGGTCACC3’ from HeLa cells. RACK1-FLAG expression construct was PCR-amplified from cDNA with Phusion polymerase (NEB) with primers CACCATGACTGAGCAGATGACCCTTCGTG TTATCACTTATCGTCGTCATCCTTGTAATCGCGTGTGCCAATGGTCACCTGC CAC and cloned into pENTR D-TOPO vector. Using Gateway LR Clonase II (Invitrogen) RACK1-FLAG was cloned into pLenti CMV Puro DEST (w118-1) (Addgene plasmid #17452) following the manufacturer’s protocol. RACK1-FLAG was expressed in RACK1 KO#1 cell line following lentiviral transduction. For lentivirus transduction, HEK293 FT cells (ThermoFisher) were co-transfected with plasmids pCMV-dR8.2 dvpr (gag-pol; Addgene #8455), pCMV-VSV-G (envelope; Addgene #8454), pAdVantage (Promega E1711) and pLenti RACK1-FLAG using Fugene HD (Promega). The lentivirus-containing media was filtered through a 0.45 μm PES filter. Following addition of 8 μg/ml protamine sulfate the supernatant was used to transduce RACK1 KO #1 cells. Cultures transduced overnight were selected with 1 µg/ml puromycin (InvivoGen) to generate a pool of stably expressing RACK1-FLAG cells. Selection was complete when untransduced RACK1 KO #1 cells had died.

### Polysome Profile Analysis

Polysome profile analyses were performed on a 10 cm dish of approximately 80%-90% confluency in 10-50% sucrose gradients containing either in 500 mM KCl, 15 mM Tris-HCl pH 7.5, 15 mM MgCl_2_, 1 mg/ml heparin (Sigma) and 100 μg/ml cycloheximide (Sigma) or 500 mM KCl; 15 mM Tris-HCl, pH 7.5; 2 mM MgCl_2_; 1 mg/ml heparin (Sigma), 2 mM puromycin as previously described by Fuchs et al. (70).

### Immunoblotting

Total protein lysate was harvested in RIPA buffer (70), and proteins separated by 12% SDS-PAGE were transferred to a nitrocellulose membrane (GE Healthcare) for 70 Vh at 4°C. Following transfer, membranes were blocked in 1% milk in PBS for 30 minutes at room temperature, washed three times in phosphate buffered saline with 0.1% (w/v) Tween 20 (PBS/T) for 10 minutes each and placed in primary antibody overnight at 4°C. Primary antibodies used were rabbit RACK1 (1:2000 dilution, Cell Signaling #4716S), FLAG-HRP (1:1000, Sigma-Aldrich, #F2555), L13a (1:1000, Cell Signaling #2765S), actin (1:1000, Sigma-Aldrich #A2066) and RPS25 (1:1000, abcam, #ab102940). Following overnight incubation, membranes were washed three times in PBS/T for 10 minutes each. For visualization of the HRP-conjugated FLAG antibody, membranes were directly imaged on a BioRad ChemiDoc XRS+. For all other antibodies, membranes were incubated in a 1:10,000 dilution of goat anti-rabbit secondary (Jackson) in 5% milk and PBS/T. Membranes were washed a final 3 times in PBS/T for 10 minutes each prior to being imaged on the BioRad ChemiDoc XRS+.

### Dual-Luciferase Assays

Bicistronic DNA constructs with *Renilla* and Firefly luciferase sequences flanked a viral IRES sequence. Viral IRESs evaluated were hepatitis C virus (HCV), cricket paralysis virus (CrPV), poliovirus (PV), and encephalomyocarditis virus (EMCV) (all gifts from Peter Sarnow, Stanford, USA). Approximately 20,000 cells of each cell line were seeded in the wells of a 96-well plate. For each construct, 100 ng DNA was transfected using lipofectamine 3000 reagent in accordance with the manufacturer’s instructions (ThermoFisher). After 24 hours, cells were lysed in 50 μl 1X passive lysis buffer (Promega) and 20 μl sample was read for 10s in a Glomax 20/20 luminometer (Promega) using the dual-luciferase assay reagent (Promega #E1910). Averages of the Firefly over *Renilla* ratio of at least three independent replicates and the standard error of the mean were calculated and plotted after normalization to wildtype cells (100%, dotted line). Following normalization to wildtype cells, statistical analysis was performed by Student’s t-test (two-tailed, unequal variance) and p-values are indicated in figure.

### Poliovirus Plaque Assays

Approximately 2 million cells were seeded into 60 mm dishes the day prior to infection. Cells were washed in PBS+ (PBS supplemented with 10 mg/ml MgCl_2_ and 10 mg/ml CaCl_2_) and 100 μl diluted poliovirus was used to infect cells for 30 minutes at 37°C, 5% CO_2_. Cells were covered with 1% bactoagar-media mixture (DMEM supplemented with 5% fetal bovine serum and 2 mM L-glutamine). After 40 hours at 37°C, 5% CO_2_, the agar layer and cells were fixed and stained for 30 minutes at RT with a crystal violet solution containing 1% formaldehyde, crystal violet and 20% ethanol. PV plaque sizes were determined by measuring the plaque diameter in pixels using ImageJ (NIH). Poliovirus plaques were counted, and average and standard error of the mean of three independent replicates were plotted. P-values were determined via Student’s t-test (two-tailed, unequal variance).

### ^35^S metabolic labeling

Wildtype, RACK1 KO #1 and #2, and RACK1-FLAG expressing cells were either mock-infected or infected with PV Mahoney at MOI = 1 (MOI = 10 for positive control) for 30 minutes at 37°C, 5% CO_2_. Cells were incubated for 3 hours and 5 minutes at 37°C, 5% CO_2_, then media was exchanged for DMEM lacking cysteine and methionine (Corning®). After starvation for 30 min, cells were metabolically labeled for 10 min with 10 μCi EasyTag™ EXPRESS^35^S Protein Labeling Mix (Perkin Elmer). Total cell lysate was harvested in RIPA buffer (70) and separated by 10% SDS-PAGE. The dried gel was exposed to a phosphor-screen (GE) and scanned using a Typhoon scanner (GE).

### Poliovirus Reporter Translation Assay

The luciferase-expressing, poliovirus-derived replicon plasmid prib(+)Luc-Wt was linearized with Mlu I (71) and *in vitro* transcribed with HiScribe T7 Quick High Yield RNA Synthesis Kit (NEB) as previously described (52). Wildtype, RACK1 KO #1 and #2, and RACK1-FLAG expressing cells were trypsinized, resuspended and gently pelleted. For each transfection reaction, approximately 3×10^6^ cells were resuspended in 500 µl of media and reverse transfected in suspension with Lipofectamine 3000 following the manufacturer’s transfection protocol for a 6-well plate. Immediately after transfection, three aliquots of 250 µl of transfected cells were mixed with 500 µl of IMDM media each. To inhibit translation, one aliquot of transfected cells was treated with 25 μg/ml cyloheximide (Sigma). In a second aliquot, PV replicon replication was inhibited with 2 mM guanidine hydrochloride. Aliquots of 150 µl were removed 3, 5, 7 and 9 h post transfection and luminescence was measured with luciferase assay reagent (Promega) using a Glomax luminometer (Promega).

## Acknowledgements

We thank Peter Sarnow (Stanford) his kind gift of viral IRES plasmids. The prib(+)Luc-Wt plasmid was a kind gift from Raul Andino (UCSF). We further thank Sangeetha Selvam and Cara Pager for discussions and for critically reading the manuscript. This work was supported by start-up funds from the University at Albany, State University of New York, and the University at Albany Faculty Research Awards Program (FRAP) (to GF).

## Author contributions

EL, CMM, and GF designed the study and wrote the paper. CMM performed RACK1 immunoblot and quantification, and 35S pulse-labeling experiment EL performed polysome and PV plaque assay experiments. EL and NP performed the dual luciferase assays, and EL and GF performed the cell-based PV-Luc replicon experiments. AJ cloned the RACK1-FLAG construct for lentivirus transduction, and ETM performed statistical analysis. All authors analyzed the data and approved the final version of the paper.

